# Stacking effects on mutation detection by T4 DNA ligation within dimeric DNA origami triangle barcodes for single-molecule nanopore analysis

**DOI:** 10.1101/2024.01.09.574918

**Authors:** Dorothy Aboagye-Mensah, Samuel Confederat, Fahad Alammar, Lekshmi Kailas, Abimbola F. Adedeji-Olulana, Alex Stopar, Allen W. Nicholson, Neil H. Thomson, Paolo Actis, Matteo Castronovo

## Abstract

Solid-state nanopores represent an emerging technology for the highly sensitive detection of biomolecular markers, but the detection of DNA point mutations is challenged by the high noise levels associated with solid-state nanopore reading. In contrast, barcoded DNA origami nanostructures can provide unique single-molecule nanopore fingerprints. In this work, we have integrated nanopore-barcoded DNA nanostructures with enzymatic DNA ligation, the latter of which is routinely involved in clinical protocols for DNA mutation detection. We designed two triangular DNA origami variants containing three elongated staples that provide strands extensions on one side that are complementary to a target sequence. Addition of the latter in solution promotes the formation of a DNA triangle dimer. Since T4 DNA ligase repairs a nick in a dsDNA segment only if there is Watson-Crick base-pairing at the nick, the two DNA triangles can be covalently linked only if the DNA sequence bridging the two triangles carries the targeted mutation. We have found striking differences between ligation detection by gel electrophoresis, AFM, and quartz capillary-based nanopores. The stacking interaction between DNA triangles is enhanced by the formation of dimers, and promote the formation of higher order nanostructure, which serve as molecular weight amplification for DNA ligation in gels. The triangle-triangle stacking dynamics presumably involves a clam-like folding mechanism, which is detectable by quartz nanopore analysis, and which hinders ligation by T4 DNA ligase. The results provide the basis for development of rapid, highly sensitive, and affordable high-throughput approaches for profiling genetic variations in point-of-care settings.

## Introduction

The development of rapid, highly sensitive, high-throughput, and affordable approaches for profiling genetic variation, including DNA point mutations in scarce sub-populations, while also avoiding information loss due to sample averaging, represents an important, as-yet unmet technological challenge.^1-3^ Such innovation is expected to have a major impact on a broad range of areas, from basic biology to food safety and personalised medicine.^2, 4, 5^

There are limitations in profiling genetic variation using current technologies. Digital(d)-PCR is a fluorescence-based approach able to provide single-molecule resolution,^6^ but has very limited multiplexing capability due to the relative scarcity of fluorescent dyes with non-overlapping spectra.^7, 8^ Multiplexed DNA point mutations analysis therefore typically involves DNA microarrays ^9^ or next-generation sequencing (NGS) technologies.^3, 10^ NGS has the advantage of allowing the untargeted analysis of genetic variants, but both technologies routinely require DNA amplification to generate sufficient DNA for analysis, which is not optimal for point-of-care applications.^1, 9, 10^ This is particularly critical for NGS analysis of suboptimal DNA samples, for which data analysis is challenged by the DNA-copying errors and bias that can occur during amplification.^11^

DNA sequencing with biological nanopores represents a new frontier in nucleic acid analysis, in which DNA molecules are detected through the analysis of single-molecule nanopore current readouts. This technology uses high-throughput, pocketsize devices,^12-14^ e.g. the MinION device from Oxford Nanopore Technology allows reading >100 kilobase DNA molecules.^15^

Solid-state nanopores (SSNP) represent an emerging alternative to biological nanopores, owing to their higher durability and lower cost, and have multiple applications in single-molecule studies.^12-14^ A quartz micropipette-based SSNP was applied to study the readout of a linear, water-soluble DNA nanostructure (“DNA beam”) comprised of a >7 kilobase ssDNA molecule hybridised to several, short ssDNA molecules, forming periodic structural repeats along the longitudinal axis of the DNA beam.^16^ Passage of the DNA beam through the nanopore produces a single-molecule, “barcoded” SSNP readout, in which each motif produces an identifiable nanopore current signature.^16^ This technology was applied to the multiplexed detection of proteins using the DNA beam as a linear nanoarray, in which a single copy of the target protein binds to one side of the beam, and a protein-specific barcode sits on the other side.^16^ An attempt to detect DNA point mutations in short DNA strands using this technology has been reported,^17^ together with alternative approaches for RNA and protein detection.^18-24^ Due to their larger diameters compared to biological nanopores, SSNPs are not yet able to accurately read DNA nucleotide sequences. Also, the DNA beams that can be resolved by SSNP with single-molecule resolution ^16, 17^ require that each target molecule captured on the beam is kept in place by a noncovalent, affinity-based, probe-target interaction.^16-18^ Since the probe-target complex is the barcode digit that encodes for target detection, accurate nanopore readout is critically dependent upon complex stability.^16-18^

To overcome this issue, in this work we have undertaken an alternative solution to achieve single-molecule DNA mutation detection using solid-state nanopores, by exploiting the unique ability of DNA nanotechnology to create barcoded nanostructures for SSNP readout. In our approach we plan to generate mutation-dependent DNA nanostructures by enzymatic DNA ligation. Ligation-based genotyping protocols have been established in clinics and involve signalling the presence of a single-base DNA mutation by enzymatic repairing of a DNA nick formed by a pair a ssDNA primers annealed to a ssDNA target.^25-27^

In this study, we designed two triangular DNA origami variants with three elongated staples forming overhang extensions on one side, as shown in the diagram in Fig. 1a. These strands are complementary to a target sequence, whose addition in solution promotes the formation of a DNA triangle dimer, as shown in Fig. 1b. Since T4 DNA ligase repairs a nick in a dsDNA segment only in the presence of Watson-Crick pairing at the nick,^28^ the two DNA triangles are covalently linked only if the DNA sequence bridging the two triangles carries the targeted mutation.

**Figure 1.**
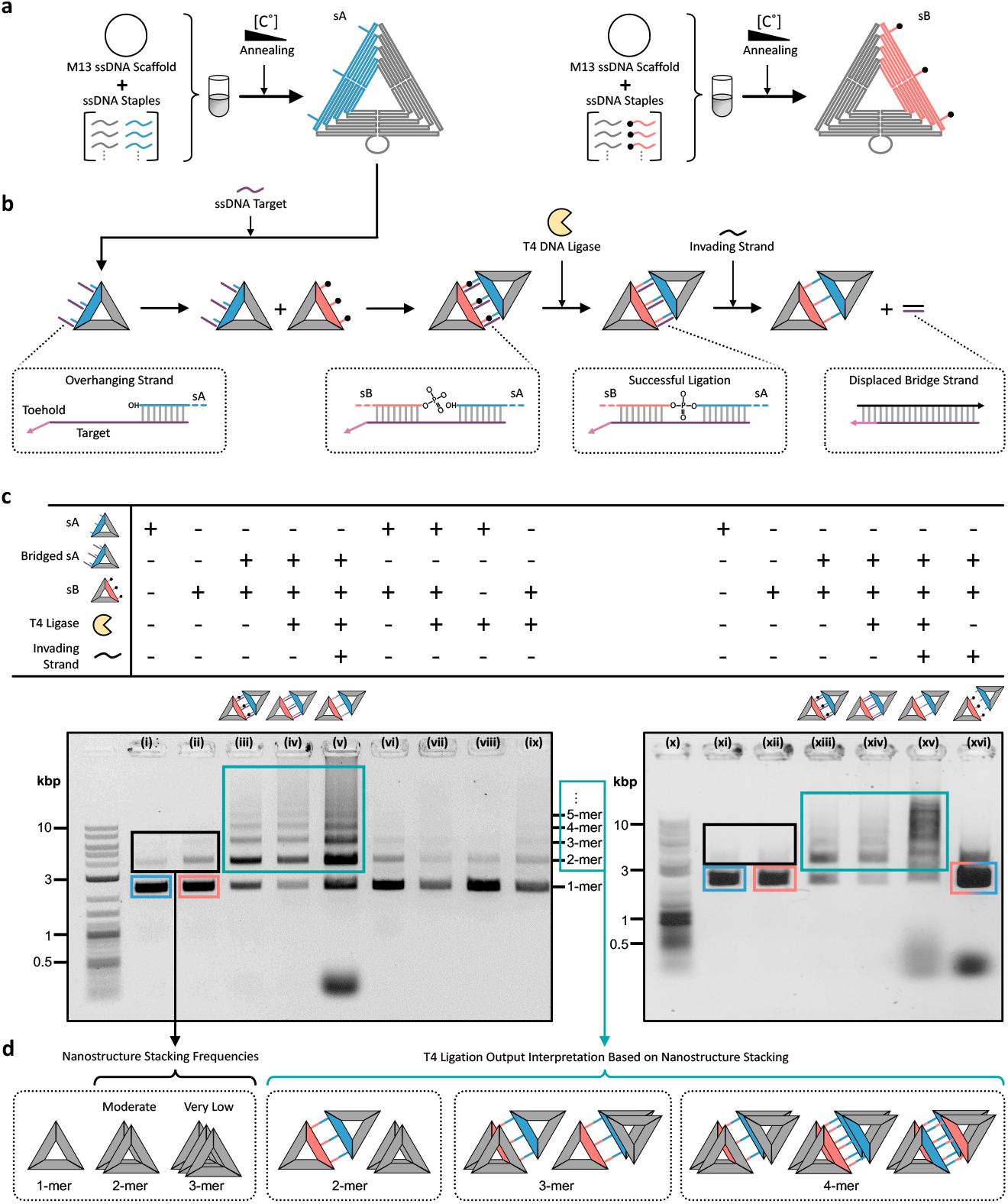
Agarose gel analysis of T4 ligation in DNA nanostructure dimers. (**a**) Schematic illustration of the self-assembly of two triangular DNA origami variants (sA and sB, respectively). For each variant, the M13 ssDNA scaffold (black circular shape) is hybridised to a structure-specific pool of short single-stranded oligodeoxynucleotides, with overhanging staple strands on different trapezoids that make up the triangle (**b**) On the left, the two variants form a dimer upon hybridization to a fully complementary ssDNA target, which form three identical dsDNA bridging links, each containing a nick in the middle. In the centre, the T4 DNA ligase repairs the nick by virtue of the phosphorylated 5’ end on the sB overhangs. On the right, a strand displacement reaction converts the three links in single strands only if the T4 ligation reaction successfully repaired the nick in the earlier step. (**c**) Gel electrophoresis characterisation of DNA triangle dimer formation and ligation (bottom). The reaction input associated with the output detected in each gel lane is shown in the truth table on top. (**d**) (right) Schematic interpretation of the dimer-dependent multimeric nanostructures shown in the gels, empirically based on the frequency of multimeric nanostructures formed by monomers (on the left).

We analysed nanostructure dimerization with agarose gel electrophoresis, atomic force microscopy (AFM) and quartz capillary nanopore reading. We found that DNA stacking interactions are enhanced by the formation of DNA triangle dimers, which promote the formation of higher order nanostructures such as DNA triangle trimer, tetramer and pentamers. This effect serves as molecular weight amplification with gel electrophoresis analysis. The triangle-triangle stacking dynamics likely involves a clam-like folding mechanism, for which the two triangles in the dimer flip between the open and the closed state. This folding induces bending of target-containing dsDNA bridges between the two triangles, which hampers the action of T4 DNA ligase, leading to a drastic reduction of the ligation efficiency.

## Results

The diagram in Fig. 1a shows the assembly of the two monomeric DNA origami triangles sA and sB involving the M13 ssDNA scaffold. Fig. 1b depicts the experimental approach to study a model ligation assay that involves DNA origami triangles serving as molecular weight amplifiers. Here, three staples of the sA DNA triangle provide identical overhang strands on trapezoid A (see Fig 1a, left), each of which can capture a model ssDNA target. The latter serve as hybridisation bridges enabling dimer formation with sB, with the latter providing identical ssDNA extensions on trapezoid B (see Fig 1a, right). The so formed dsDNA segments, comprised in the cavity between sA and sB, provide substrates for T4 ligation. Finally, a toehold strand displacement reaction acting on the target sequence allows for assessing ligation efficiency, in that each yielded copy of DNA triangle dimer signals the occurrence of at least one successful ligation event in the cavity.

### Agarose gel analysis of enzymatic ligation within triangle dimers

Figure 1c (bottom) shows agarose gel electropherograms of the results obtained by varying the inputs or the reaction steps, as shown in the diagram above. The gel bands in lanes i and xi are the sA monomer, whereas the bands in lanes ii and xii are the sB variant. In both cases, the main band at the bottom indicates successful formation of the monomeric nanostructure, whereas the two less pronounced gel bands of slower mobilities (framed in black) suggest the presence of sA or sB dimers trimers. In the presence of the ssDNA target (lanes iii and xiii), multimeric structures of sA and sB are present that migrate around or above the 10 kbp reference, including a greater yield for dimers and trimers. T4 DNA ligase treatment has negligible effects on the electrophoretic profiles (see lanes iv and xiv), whereas the strand displacement reaction following ligation increases the amounts of higher order multimeric structures output (see lanes v and xv). Strand displacement in absence of prior ligation suppresses such higher order structures and promotes nanostructure monomers and dimers as the main outputs (see lane xvi). Further control experiments involving both sA and sB in the absence of the ssDNA target show that the action of ligase reduces the background of multimeric nanostructures (see lanes vi and vii, respectively). Such background is apparently enhanced in experiments involving a single nanostructure input (see lanes viii and ix, respectively). Overall, the higher-order gel bands (framed in dark cyan) match well across the board and allow the resolution of different multimeric forms, from triangle monomers to pentamers, with single-triangle increments in molecular weight.

### AFM characterisation of triangle dimers

Figure 2a shows topographic AFM images of the sA triangle monomer (left panel) and the sA-sB dimerization product (right three panels) adsorbed on mica. Of note, nanostructure monomers are predominant in the ligation. To quantitatively analyse monomer and dimer copy numbers, we assume that the presence of sA-sB dimer requires visual evidence of at least one connecting strand between the edges of two triangles. Any other configuration presenting two or more DNA triangles placed side-by-side not showing such a link are deemed to comprise monomers only. For example, only one sA-sB dimer is evident across the example AFM micrographs, as highlighted in the diagram in Fig 2b. The built-in helical loop protruding from the triangle could be confused as a connection between triangles, but can be discarded by considering the nanostructure symmetry. In addition to fully formed DNA triangles, damaged nanostructures also are visible on the right of Fig 2a, as they lack a regular triangular shape. The histogram in Fig. 2c shows the monomer vs. dimer counts in each of a set of 10 AFM images, indicating the paucity of DNA triangle dimers in our samples. The AFM analysis, therefore, does not provide structural information on the formation of triangle dimers and higher order nanostructures, the lack of which in AFM images could reflect the effect of structural alterations following surface adsorption.

**Figure 2.**
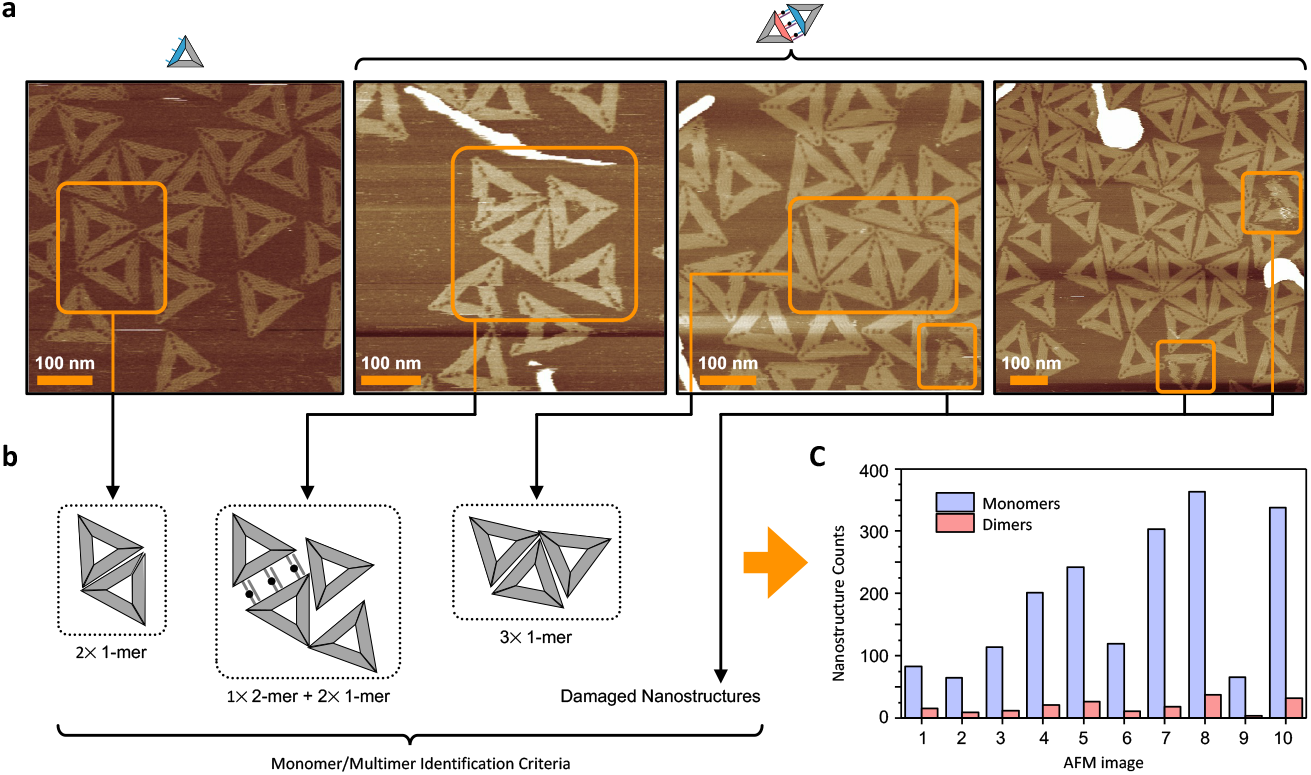
AFM characterisation of DNA nanostructures. (**a**) AFM topographic images showing self-assembled DNA nanostructures, such as monomers (left) and the output of DNA triangle dimerization (right). The scale bars are 100 nm. (**b**) Schematic representation of three groups of triangular nanostructures identified in the AFM images in (a) (highlighted with orange frames), respectively comprising two adjacent monomers, a dimer comprising three DNA links near two monomers, and three adjacent monomers. Damaged structures are defined as those lacking a regular shape or presenting missing parts, including any artefact attributed to the imaging process. (**c**) Histogram of DNA nanostructure counts (monomer in blue, and dimer in red) from 10 topographic AFM images according to the classification schematically presented in b.

### Nanopore translocation analysis of triangle dimers

Nanopore translocation studies were conducted using glass nanopipettes having a nanopore diameter of 160 nm (data not shown), which is near the nominal width of the DNA triangle monomer. Here, if an electric field is established between the electrode inside the nanopipette (containing inner electrolyte) and the electrode in the outer electrolyte (bulk electrolyte solution in the petri dish) the voltage gradient triggers DNA nanostructure translocation from the nanopipette into the outer electrolyte, as schematically represented in Fig. 3a (left). As described in the methods section, the inner/outer electrolyte consisted of 0.1 M KCl, with a molecular crowding agent (50% w/v of 35 kDa PEG) added to the outer solution to enhance the translocation signal.^29, 30^ Each translocation event produces a negative peak in the ionic current recording, which is characterised by two main parameters: the dwell time (*i*.*e*., the duration of the current fluctuation) and the peak current maxima (*i*.*e*., the module of maximum fluctuation associated with the translocation event with respect to the current baseline), as summarised in Fig. 3a (right).

**Figure 3.**
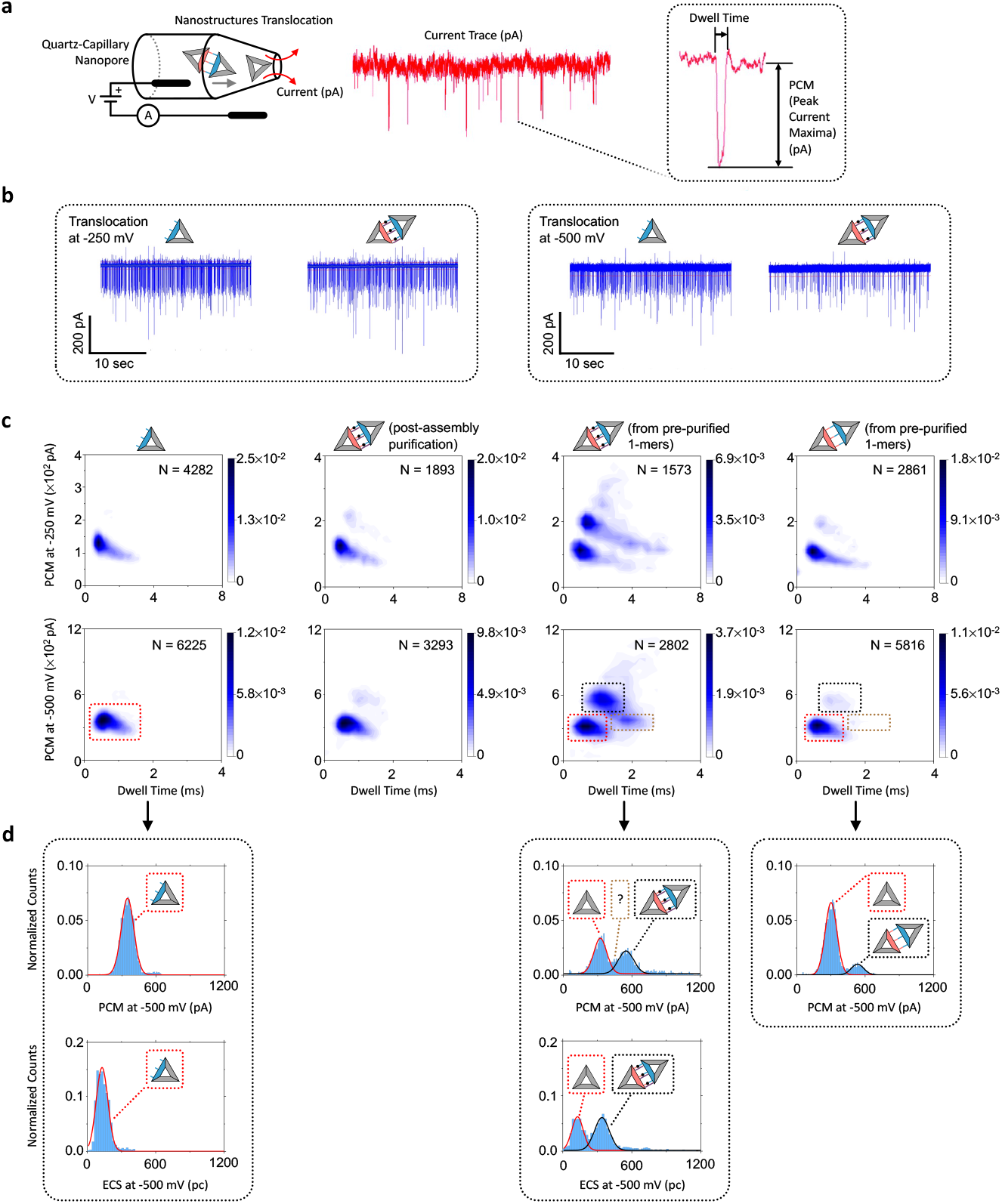
Nanopipette analysis of DNA nanostructures. (**a**) Schematic illustration of the nanopipette setup. Ion current trace showing translocation events (red trace) and a zoom-in of a single ionic current peak showing the peak current maxima (PCM), the drop in current, and the dwell time. (**b**) Exemplar ion current traces for DNA triangle monomer and dimer at -250 mV (left) and -500 mV (right). (**c**) Density scatter plots of PCM as a function of dwell time. All nanopipette experiments were carried out at -250 mV (top panels) and -500 mV (bottom panels). In each plot, the total number of detected events N is shown. (**d**) The top panels show the distribution of translocation event counts (at -500 mV) as a function of PCM corresponding to the scatter plots above. The bottom panels show the same translocation event counts as a function ECS. The gaussian fits corresponding to monomers and dimers are denoted with red and black solid line, respectively.

Representative ionic current traces of DNA triangle monomer (sA) translocation and of the product of a dimerization reaction obtained at -250 mV and -500 mV, respectively, are shown in Fig. 3b (at overall constant M13 concentration, 500 pM). Several conductive peaks associated with ionic current modulation can be observed in each current trace.^29^ Comparing the current trace of the sA monomer and the sA-sB product at both voltages (-250 mV and -500 mV) in Fig. 3b, it is observed that the ratio of low *vs*. high amplitudes is essentially constant. Nevertheless, the translocation frequency is higher with the sA monomers at both applied voltages, and this is potentially caused by an increased electrophoretic mobility of smaller structures as it has been observed before also with gold nanoparticles.^31^

The top panels in Fig. 3c show heat map plots depicting the distribution of translocation events in terms of peak current maxima (PCM) *vs*. translocation dwell time, measured at an applied voltage of -250 mV for (i) the sA monomeric structure as reference (left); the dimer formed through hybridisation, produced in two ways, *i*.*e*., (ii) formation of dimers from non-purified monomers, followed by purification and (iii) formation of dimers from purified monomers; and finally (iv) ligated dimers formed from purified monomers, subsequently treated with strand-displacement. The monomeric variant provides a background cluster with PCM slightly above 100 pA and an average dwell time of nearly 1 msec that is visible with all samples. The dimeric form shows an additional cluster at a PCM at ∼200 pA, which however is significant only for the dimerisation output generated from purified sA and sB monomers (centre-right). Surprisingly, this additional cluster is similarly diminished with post-sA-sB-assembly purification (centre-left), or by sA-sB ligation followed by strand displacement (on the right). Similar outcomes are obtained at an applied voltage of -500 mV as displayed at the bottom of Fig. 3c. Whereas the average PCM is nearly 2-fold higher, the nanostructure transit is faster as evidenced by a slightly lower average dwell time. Of note, at -500 mV the sA-sB output obtained from purified monomers shows a third cluster (highlighted by the brown box), associated with PCM values in line with the monomeric form and approximately twice the average dwell time (*i*.*e*., nearly 2 msec). Focusing on these latter samples, Fig. 3d shows on top the renormalised PCM distributions (based on the renormalisation factors N in Fig. 3c) and their gaussian fits. The reference sA monomer gives a single peak, whereas the sA-sB output show an additional yet smaller peak, which decreases after ligation and strand displacement. For the sA-sB samples, the latter peak includes values associated with the third cluster mentioned above, which does not resemble monomer transit events. Since, the amplitude/shape of the current peak with respect to the baseline and its duration can vary depending on nanostructure shape or orientation during transit, we integrated the current trace of each translocation event to obtain the distributions of equivalent charge surplus (ECS), which is less likely to depend on nanostructure geometry,^29^ and which is shown at the bottom in Fig. 3d. Interestingly, the ECS analysis for the sA-sB output gives two distinct gaussian distributions of similar amplitude associated, respectively, with the electric charge of the monomer and the dimer. This result indicates that, the clusters highlighted by the black and brown boxes in Fig. 3c are possibly associated with DNA triangle dimers translocating with different structural/conformational modalities (see Discussion).

## Discussion

This study has investigated the integration of DNA nanotechnology, DNA ligation, and nanopipette current analysis, towards the development of an amplification-free scheme for DNA mutation detection. The key step in the detection scheme involves the ligation-dependent formation of a DNA origami triangle dimer to yield a unique nanopore fingerprint. The investigation primarily focused on DNA nanostructure dimer formation and detection by nanopipette analysis. In our exemplar approach, to balance detection sensitivity and stability of the DNA nanostructure dimer, the stability of the latter reflected the ligation of three copies of a ssDNA target sequence (see Fig 1a). To adapt the DNA mutation detection-by-ligation scheme to our combined DNA nanotechnology and solid-state nanopore-based approach, a strand-displacement reaction was designed to output sA-sB dimers only if a successful DNA ligation reaction occurs between the flanking DNA triangles sA and sB (see Fig 1b). Furthermore, control experiments were performed to model the outcome of lack of DNA mutation detection (see the table in Fig. 1c).

Agarose gel analysis shows that both sA and sB monomers tend to form homodimers (with varying yields) as well as homotrimers, in significantly lower amounts. Otherwise, in all situations that promote sA-sB dimer formation, we noticed the presence of higher-order multimeric structures in the form of a ladder, with gel band intensity being inversely proportional to molecular weight. The dissociation (or lack of association) of the sA-sB dimer inhibits multimeric structure formation, which are therefore nanostructure dimer- and therefore target-specific.

Based on the self-association tendency of the DNA triangle monomer, the diagram on the centre-right in Fig. 1d depicts our interpretation of multimeric structure formation (including dimer, trimer, and tetramer), as based on a prevalent stacking of dimeric triangles.

Furthermore, the significant monomeric background in lane v in the gel in Fig. 1c indicates T4 DNA ligation does not reach completion despite the reaction being carried out in saturating enzyme reaction conditions. This result is consistent with previous studies of the action of T4 DNA ligase on DNA origami nanostructures, reporting efficiency values near 50%.^32, 33^

For the AFM analysis we focused on the formation of nanostructure dimers dependent upon probe-target hybridisation. The AFM image analysis, however, reveals that the fraction of DNA triangle dimers (presenting at least one dsDNA bridge) deposited on mica is significantly lower than the fraction of DNA dimers detected by agarose gel electrophoresis. Furthermore, no multimeric structures were observed. Our interpretation is that DNA triangle monomers are more efficiently adsorbed on mica, as compared with dimers or multimers, reflecting their greater roto-translational degrees of freedom.^34^ Furthermore, multimeric forms held together by DNA stacking, would be easily destabilised by surface charges and enhanced surface diffusion after mica adsorption.^35^

Since nanopore analysis allows reading time-dependent current traces that depend on the translocation behaviour of single DNA nanostructures transiting through the pore, it can provide complementary information on the structural properties of nanostructure dimers, as well as other multimeric forms in solution. The data in Fig. 3c confirm the presence of the monomeric background observed with gel electrophoresis. It also reveals the presence of the dimers when sA-sB dimer formation is promoted, which, in contrast to our in-bulk gel analysis, is dependent upon sample processing.

Interestingly, dimers are formed from pre-purified monomers in higher yields than dimers that are purified after sA-sB formation. Furthermore, the association of pre-purified sA and sB monomers produces a more complex nanopore readout, with sporadic current peak events of much higher intensity, and duration than the reference values for monomer and dimer. The results obtained in a previous study focusing on the nanopore characterisation of the multimeric association of DNA origami squares^29^ suggest that these sporadic events reflect the multimeric structures detected by gel electrophoresis.

The absence of such high-intensity nanopore current peaks from samples purified after sA-sB hybridisation suggests that the presence of ssDNA staples in solution disrupt dimer formation and relatively weak stacking interactions present in the multimeric nanostructures. Furthermore, T4 DNA ligase treatment followed by strand displacement (i) suppresses the sporadic, high-intensity nanopore current peaks; (ii) significantly reduces the frequency of dimeric events; and (iii) consistently promotes monomeric events (see Fig. 3c). Taken together these results show that ligation efficiency is considerably lower than 50%. Furthermore, the comparison of the gel and nanopore results suggests that DNA migration in the gel matrix preserves multimeric DNA nanostructures, even after the action of T4 DNA ligase followed by the displacement of the ssDNA target connecting sA and sB. In contrast, the relatively weak stacking interactions holding multimers in this configuration are challenged by the shear stress in the quartz microcapillary environment.

The PCM and ECS analyses allowed detection of two distinct nanopore transit modalities for the DNA triangle dimer. In the main cluster, the dwell time is < 1 msec and the PCM > 500 pA, while in the minor cluster the dwell time is nearly 2 msec and PCM < 500 pA. The diagram in Fig. 4a depicts our interpretation of the coexistence of these to dimer-related populations, while the diagram in Fig. 4b presents a qualitative scoring of these observations. The first dimer-related (more populated) cluster corresponds to a self-stacking sA-sB dimer, enabled by the presence of a nick in the dsDNA segments connecting the two triangular units. This form would have the same longitudinal width of a DNA triangle monomer, but higher mechanical rigidity. The second (less populated) cluster corresponds to an open sA-sB dimer, which, compared to the monomer, would have twice its longitudinal width and similar mechanical rigidity. A comparison of the the different cluster populations (see Fig 3c right) indicates that the self-stacking form is more frequent in solution, likely due to higher stability. Of note, in this latter configuration, the dsDNA segments connecting sA and sB are bent and therefore structurally unavailable to bind T4 DNA ligase. This would explain the relatively low enzymatic efficiency, as evidenced by the PCM distribution shown on the right in Fig 3d.

**Figure 4.**
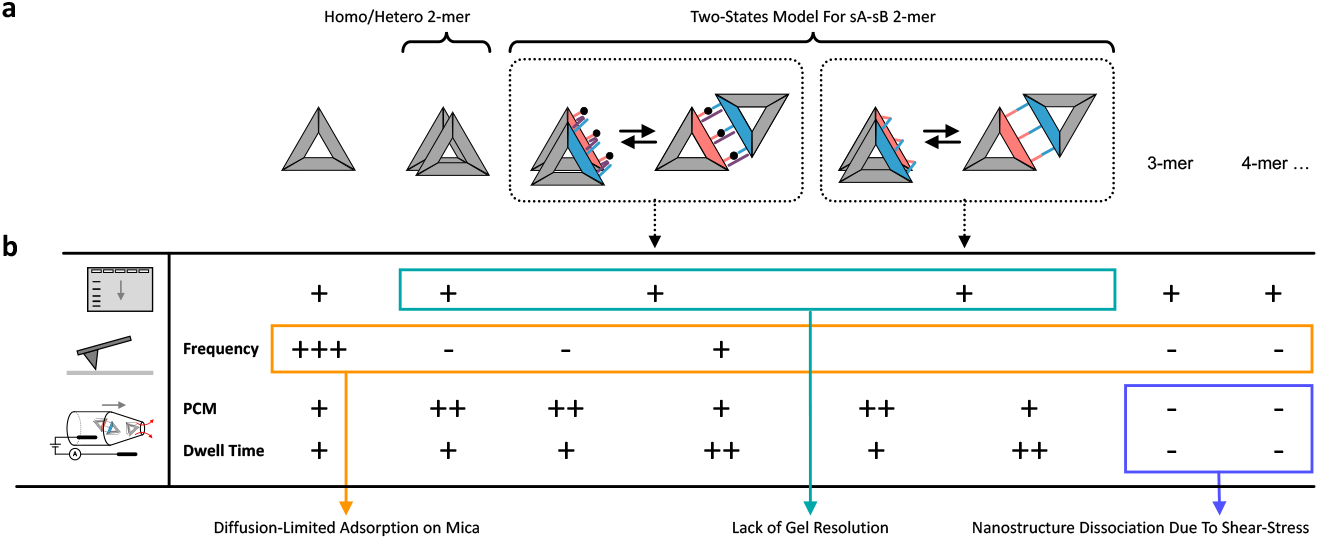
Empirical analysis of structural factors affecting DNA ligation in a DNA triangle dimer. (**a**) Diagrams of different DNA nanostructure forms. In particular, a two-state model is shown for the sA-sB dimer, in which the DNA triangles are either stacked or planar. (**b**) The table qualitatively scores the frequency of each structure shown in the diagram in (a) as a function of the adopted analytical approach. Gel electrophoresis detects the formation of multimeric structures likely formed by stacking interactions between DNA nanostructures (hence a “+” is shown for all structures) but cannot discriminate the two-states that we predict for the sA-sB dimer. AFM imaging mainly detects monomeric DNA triangles, likely due to diffusion limited adsorption on mica and/or surface stabilisation effects. Nanopipette translocation analysis successfully detects dimeric nanostructures. Here, dimers are associated with either high PCM (++) and short dwell (+) time, or low PCM (+) and long dwell time (++), thus allowing resolving the two-state model for the dimeric form (extended or folded). The nanopipette analysis cannot detect multimeric nanostructures observed with gel electrophoresis, which in the nanopipette are likely dissociated by sheer stress during nanopore translocation.

In summary, we have explored a DNA nanotechnology approach that integrates a classical DNA-mutation-by-ligation assay with solid-state nanopore detection. In this approach, DNA mutation detection outputs a stable DNA origami nanostructure dimer. We have therefore investigated DNA triangle dimer formation and the action of T4 DNA ligase that functions in the localised environment. We have found striking differences between the results obtained with gel electrophoresis, AFM, and quartz capillary-based nanopores. Namely, with gel electrophoresis we detected the unexpected formation of dimer-specific, higher-order, multimeric structures likely formed by stacking interactions between DNA triangles. In contrast, with single-molecule AFM or nanopore analysis, such multimeric structures are challenged by the presence of surface effects on mica (affecting AFM imaging) or confinement in quartz microcapillary (causing higher shear stress during nanopore translocation). Finally, we have found that the action of T4 DNA ligase is inhibited by the presence of triangle-triangle stacking effects. Overall, these results will serve to inform the design of novel nanopore-based DNA mutation-sensing schemes.

## Materials and Methods

### DNA origami nanostructure design

The DNA origami structures were designed using CaDNAno version 1.0 software (CaDNAno, n.d.), following the triangular nanostructure designs published by Rothemund.^36^ For this study, two triangular DNA origami variants were designed. Both triangles had three elongated staples forming part of the main body of the nanostructure, forming overhang strands protruding from the longer parallel side of one of the trapezoid sub-structures of the triangle. One had its overhang strands protruding from trapezoid A and is hereafter termed “sA”, the other had its overhang strands protruding from trapezoid B is hereafter termed “sB”. Each overhang strand was 20 bases long and they were positioned in the middle and the two corners of the longer parallel side of the trapezoids. We designed protruding strands from sB that were phosphorylated on the 5’end to enable the action of T4 DNA ligase. We also designed a target strand (37 bases long) complementary to the 15 bases of the overhang sequences of the two DNA triangles (sA and sB). In addition, a 7 base long sequence, serving as toehold domain, was added to the 5’ end of the target strand DNA triangles sA and sB come into proximity and alignment when the target strands hybridise with the overhang strands from both DNA triangles, thus forming a nicked dsDNA stem on which the T4 DNA ligase can act. Furthermore, we designed a 37-base long, invading ssDNA strand fully complementary to the target sequence. The hybridisation between the latter and the free ssDNA toehold domain of the target strand promotes a branch migration reaction leading to irreversible target displacement from the sA-sB dimeric complex.^37, 38^ The 20-base overhang strand is in red, whereas the DNA sequence in black denotes the original staple located inside the DNA origami. For the target and invading strands, the DNA sequence in blue denotes the toehold domain.

Elongated staples forming overhang strands on triangle sA:

5’-

GAGAATCGAATCGGCTGTCTTTCCAAGGCTTATCCGGTATGCAAACACAACATCCCACAG -3’

5’-TTTATCCTGAATCTTACCAACGCTAACGAGCGGCAAACACAACATCCCACAG -3’

5’-CAATCAATAACCCACAAGAATTGACAGAGAGAATAACATAGCAAACACAACATCCCACAG -3’

Elongated staples forming overhang strands on triangle sB:

Phos-5’-GCAAACACAACATCCCACAGTCTTGACAAGAACCGGTCGCCTGATAAATTGTTCGGAACG – 3’

Phos-5’-GCAAACACAACATCCCACAGACTTTAATCATTGTGATCCAATACTGCGGAATGCTTTAAA – 3’

Phos-5’-GCAAACACAACATCCCACAGTTCATCAGTTGAGATTGCATAGTAAGAGCAACTCGCGTTT TAA – 3’

Target Strand with toehold domain:

5’-ATGGAGAGGATGTTGTGTTTGC CTGTGGGATGTTGTG – 3’

Invading strand with toehold domain:

5’-CACAACATCCCACAGGCAAACACAACATCCTCTCCAT-3’

### DNA origami nanostructure preparation

The M13mp18 single-stranded DNA scaffold was purchased from New England Biolabs (NEB) at an initial concentration of 250 µg/mL. The staple strands were purchased from Sigma-Aldrich and resuspended in distilled water to a final concentration of 100 µM. The target and invading strands were purchased from Integrated DNA Technologies (IDT) and resuspended in distilled water to a final concentration of 100 µM.

The DNA origami objects were self-assembled in one-pot reaction mixtures, consisting of a circular ssDNA scaffold (10 nM) and staples (50 nM) in a final volume of 200 µL in a buffer containing 40 mM Tris, 12.5 mM MgCl_2_, pH 8.5. Self-assembly of the DNA origami object was initiated by heating the reaction mixture at 95 °C for 5 minutes, then decreasing the temperature from 95 ·C to 20 ·C at a rate of 1 °C per minute in a thermal cycler (Bioer GeneTouch thermal cycler). The self-assembled nanostructures were purified using Sephacryl S400 (GE Healthcare) size-exclusion columns to remove the staple strands in excess. Subsequently, the DNA origami nanostructures were eluted in the self-assembly buffer (40 mM Tris, 12.5mM MgCl_2_, pH 8.5) and the concentration was measured using a NanoDrop 2000c Spectrophotometer (Thermo Fisher Scientific Inchinnan, UK). The purified Sa nanostructure was first hybridised to the complementary target strands in the ratio 1:5, respectively, at a final volume of 50 µL, in 40 mM Tris, 12.5 mM MgCl_2_, pH 8.5. The solution was then incubated at 20 ·C on a heat block for 1 hour. The target-capturing sA nanostructure was then hybridised to sB in a 1:1 stoichiometry at a final volume of 50 µL in 40 mM Tris, 12.5 mM MgCl_2_, pH 8.5, then incubated at 20 ·C for 1 hour on a heat block. Subsequently, the nanostructures were subjected to a T4 ligation reaction containing 5 nM of sA-sB complex and 10 Weiss units of T4 DNA ligase in a buffer containing 40 mM Tris, 10 mM MgCl_2_, 10 mM DTT, 0.5 mM ATP, pH 7.8, in a final volume of 35 µL. The ligation reaction was allowed to proceed for 2 hours at 25 ·C using a heat block for incubation. Subsequently, the strand displacement reaction was carried out at 20 ·C for 1 hour by incubating the sA-sB complex at 4.8 nM and the invading strand (at a stochiometric ratio of 1:5) in a 30 µL solution comprising 40 mM Tris and 12.5 mM MgCl_2_, pH 8.5.

### Gel electrophoresis analysis

To detect the formation of nanostructure sA, sB and any higher order complexes, we utilised agarose gel electrophoresis (AGE) to sort out different nanostructures according to their molecular weight. To this end, 0.7% agarose gel was cast and stored at 4 ·C overnight, followed by the loading of approx. 200 ng of DNA nanostructure together with 1X orange loading dye. The DNA was electrophoresed in 1X TAE buffer (pH 8) at a constant voltage of 60 V for 3 hours on ice. A 1kbp DNA marker (GeneRuler 1 kb plus Thermo Fisher Scientific) was used to estimate the molecular weight of the DNA nanostructures. For the post-staining of the gels, the gels were either stained with 0.5 µg/mL of ethidium bromide solution or 1X SYBR safe (Thermo Fisher Scientific) and followed by de-staining in Milli-Q water for 15 minutes. Lastly, the gels were visualised using Molecular imager Chemi-DOC MP imaging system (Bio-Rad) and analysed using ImageJ software version 2.0.

### AFM imaging

For AFM imaging experiments, DNA samples were diluted to a final concentration of 0.5 nM in a solution comprising 40 mM Tris, 12.5mM MgCl_2_ at pH 8.5. 30 µL of DNA sample was deposited onto a freshly cleaved mica disc glued on top of a 15 mm diameter metal sample disc, and then incubated at room temperature for approx. 30 minutes in a covered petri dish. The mica surface was then rinsed gently with filtered Milli-Q water about ten times and then placed on the AFM sample stage. An additional 100 µL of the self-assembly buffer (40 mM Tris and 12.5 mM MgCl_2_ at pH 8.5) was placed onto the mica disc to allow enough solution to cover the sample surface and prevent sample drying during imaging. The AFM imaging was carried out in Tapping mode in liquid on a Bruker Dimension FastScan Bio AFM (at the Bragg Centre for Materials Research, Leeds) with NanoScope 9.4 software, using FastScan-D from Bruker (resonant frequency of ∼110 kHz and spring constant of 0.25 N/m) and scan rate ranging between 1 Hz – 1.5 Hz. Between 5 and 10 AFM images were acquired at 1024 x 1024 pixel resolution for each triangular DNA origami variant from different areas per sample. The obtained topographic images were processed with 1^st^ order flattening and analysed using NanoScope Analysis software version 1.9. (Software, n.d.).

### Nanopore translocation measurements

The glass nanopipettes used were fabricated from quartz capillaries with filament (QF100-50-7.5, World Precision instrument, UK) using a P2000 laser puller (Sutter instrument, 2020). The quartz capillaries are 7.5 cm long with an outer diameter of 1.0 mm and an inner diameter of 0.5 mm., while the formed nanopipettes had a pore diameter ranging from 80-100 nm.^19^ The translocation experiments were carried out by filling the nanopipette with the translocation buffer (100 mM KCl, 0.01% Triton-X, 10 mM Tris, 1 mM EDTA pH 8.0) containing the DNA origami sample at a concentration of 500 pM. The nanopipette was then immersed in a 100 mM KCl bath with the addition of 50% (w/v) polyethylene glycol (PEG) 35 kDa (ultrapure grade, Sigma Aldrich). A working electrode consisting of Ag/AgCl wire (0.25 mm diameter, GoodFellow, Huntingdon, UK) was inserted in the nanopipette barrel. In addition, a second Ag/AgCl wire was immersed in the bath and acted as the reference electrode. The DNA origami nanostructures were translocated from inside the nanopipette into the external bath by applying a negative potential to the working electrode placed inside the nanopipette with respect to the reference electrode in the bath. The transiting of DNA origami nanostructures with a size near the diameter of the nanopipette tip orifice causes a temporary physical blockage, which hinders the free-flowing electrolyte ions, thus resulting in a drop of ionic current. The ion current was recorded with a MultiClamp 700B patch-clamp amplifier (Molecular Devices, San Jose, CA, USA) in voltage-clamp mode. Data were acquired at a 100 kHz sampling rate with a 20 kHz lowpass filter using the pClamp10 software (Molecular Devices). The ion current traces were further analysed with the MATLAB script Transanlyser, developed by others.^39^ The obtained translocation events were analysed by applying a 7-sigma threshold level from the baseline, and only the events with peak current above the threshold were considered successful translocation events. All data were further analysed and plot using Origin 2019b, see Fig. 3.

## Acknowledgements

DA-M, AFA-O, AWN and MC acknowledge funding from the European Union’s Horizon 2020 research and innovation program under the Marie Sk1odowska-Curie MSCA-RISE grant agreement no. 645684. S.C. and P.A. acknowledge funding from the European Union’s Horizon 2020 research and innovation program under the Marie Sk1odowska-Curie MSCA-ITN grant agreement no. 812398, through the single-entity nanoelectrochemistry, SENTINEL project. LK was supported by EPSRC Strategic Equipment Scheme grant (EP/R043337/1) as Experimental Officer for the Leeds AFM facility. The Bruker FastScan BioAFM was acquired via a Wellcome Trust equipment grant (101497/Z/13/Z). PA acknowledges funding from the Engineering and Physical Sciences Research Council UK (EPSRC) Healthcare Technologies for the grant EP/W004933/1.

